# The PAAR5 Protein of a Polarly-Localized Type VI Secretion System (T6SS-5) is Required for the Formation of Multinucleated Cells (MNCs) and Virulence by *Burkholderia pseudomallei*

**DOI:** 10.1101/2023.02.09.527577

**Authors:** Christopher T. French, Philip Bulterys, Javier Ceja-Navarro, David Deshazer, Kenneth Ng

## Abstract

The *Burkholderia pseudomallei* complex (Bpc) includes *B. pseudomallei, B. mallei* and *B. thailandensis*. These species share conserved virulence determinants that facilitate survival in mammalian cells and can spread from cell to cell by a unique mechanism involving fusion of plasma membranes. The activity of a contractile type VI secretion system, T6SS-5, is a central requirement. Using fluorescence confocal microscopy, we found localization and dynamic turnover of fluorescently-labeled T6SS-5 components at the forward pole of *Burkholderia* residing at the ends of actin protrusions. We identified the proline-alanine-alanine-arginine repeat protein of T6SS-5 (PAAR5), which forms the heteromeric tip of the T6SS-5 apparatus along with VgrG5. Mutational analysis revealed a unique N-terminal extension (NTE) of PAAR5 that is indispensable for cell fusion. Deletion of *paar5* allowed us to uncouple fusogenic activity from the functionality of T6SS-5 for exploring the role of cell fusion in pathogenesis. *B. pseudomallei* Δ*paar5* deletion mutants retained a functional T6SS-5 apparatus and the ability to secrete the Hcp5 protein. In cellular and animal infection models, Δ*paar5* mutants mirrored the phenotype of a T6SS-5-defective Δ*vgrG5* strain, being defective for cell fusion and avirulent in hamsters. These results demonstrate concordance between the fusogenic and *in vivo* virulence phenotypes, suggesting that T6SS-5-mediated cell fusion may be a central feature of *B. pseudomallei* pathogenesis and not an *in vitro* artifact.

## INTRODUCTION

*Burkholderia pseudomallei* (*Bp*) is a Gram-negative bacterium that inhabits moist, tropical soils and causes melioidosis in humans and animals. According to estimates, *Bp* is endemic in more than 79 countries with a combined population of over 3 billion, and is responsible for 165,000 annual human infections with a mortality rate near 50% (1). *B mallei* (*Bm*), which causes glanders, is a derivative of *Bp* that has adapted to a lifestyle of obligate parasitism in mammals. The genome of *Bm* has undergone a considerable reduction in size, losing loci needed for survival in the environment while retaining those for survival in a mammalian host (2, 3). Although glanders is primarily a disease of equids, severe and life-threatening human infections do occur (2, 4). *Bm* and *Bp* are classified by the U.S. Centers for Disease Control and Prevention as Tier-1 Select Agents due to their high infectivity, severity of disease, broad antimicrobial resistance, and historical use in biological warfare (5, 6). A third member of the *Burkholderia pseudomallei* complex species (Bpc spp.), *B. thailandensis* (*Bt*), shares conserved virulence mechanisms with *Bm* and *Bp*, although it is less virulent and rarely associated with disease (7–10).

*Bm, Bp* and *Bt* can survive intracellularly (11–17). Once inside, bacteria escape from membrane-bound phagosomes, replicate in the cytoplasm and spread from cell to cell (4). Escape from phagosomes is facilitated by the conserved Bsa type III secretion system (T3SS_Bsa_), which in *Bp* is also designated as T3SS-3 (12, 18). Bacterial motility is a requirement for cell-cell spread and can be provided by BimA-dependent actin polymerization or the Fla2 flagellar system, if present (12, 19–21). Intercellular spread is mediated by a type VI secretion system (T6SS-5) that catalyzes membrane fusion between adjacent cells, creating a portal for direct passage of bacteria into the cytosol of neighboring cells (12, 22). Repeated fusion events give rise to large multinucleated cells (MNCs), which is fundamentally different from multinucleated giant cell (MNGC) formation observed in mycobacterial infections or other granulomatous diseases. In those settings, MNGCs develop by cytokine-mediated activation of a cell fusion program present in monocytes and macrophages, which can be reproduced *in vitro* by adding cytokines alone (23). In contrast, membrane fusion following infection by *Burkholderia* appears to be bacterially-mediated, resulting from firing of T6SS-5 proteins across the bacterial envelope and through the plasma membranes of the infected cells, causing local deformation and merging of apposing membrane bilayers (12, 22). While cell fusion is critical for cell-cell spread and survival *in vitro*, MNCs have only occasionally been reported in tissues from infected animals and human melioidosis patients (19, 22, 24–27). *Bt* is a suitable surrogate for *in vitro* studies of *Burkholderia-mediated* cell fusion. *Bt* contains a highly related T6SS-5, its *vgrG5* spike protein locus is functionally interchangeable with *Bp* and it can be safely manipulated in BSL-2 (22).

Encoded by clusters of 15-20 genes, T6SSs are multicomponent contractile nanomachines that resemble the injection devices of tailed bacteriophages, allowing bacteria to translocate effector proteins into target cells (28–31). According to current views, the base and anchoring components of the apparatus are assembled across the inner and outer bacterial membranes. A contractile sheath of TssB/C components is assembled in the cytoplasm surrounding an inner tube composed of Hcp subunits. A feature of the T6SS is a cell-piercing tip complex composed of trimeric Val-Gly-Arg (VgrG) repeat proteins (28, 32) and a Pro-Ala-Ala-Arg (PAAR) repeat protein that caps the VgrG spike (33, 34). VgrG and PAAR proteins may contain extended domains which may bind and translocate additional effectors with activities in target cells (32, 33, 35–43). The *a la carte* nature of VgrG, PAAR and their associated effector proteins and domains confers a diverse range of functionalities to T6SSs. Firing of the apparatus correlates with sheath contraction and ejection of the Hcp tube and the associated VgrG/PAAR tip complex. The ClpV AAA-type ATPase recognizes and disassembles the contracted TssB/C sheath, and subunits are recycled for subsequent rounds of activity (44).

The genome of *Bp* encodes six distinct T6SSs, five of which are differentially represented in *Bm* and *Bt*. As the nomenclature for *Burkholderia* T6SS loci varies among prior studies, we number the individual T6SS gene clusters corresponding to their chromosomal coordinates in *Bp*. Accordingly, we refer to the system that is associated with virulence in animals as T6SS-5 (22, 26, 45–48), although earlier studies refer to the same system as T6SS-1 (12, 26, 27, 49, 50). T6SS-5 is required for MNC formation and virulence, and is conserved in *Bp*, *Bt* and *Bm*. Elucidating the mechanism of T6SS-5-mediated membrane fusion has been problematic, as has been deciphering the relationship between fusion and virulence. This is confounded by an inability to separate membrane fusion from other functions of T6SS-5, since mutations that affect fusion typically render the T6SS apparatus functionally inoperative or unable to assemble. This is further complicated by the multiple T6SS gene clusters of Bpc spp., and the numerous *vgrG* and *paar* loci within these clusters and at remote sites in the genome. In this report, we describe a model of cell fusion mediated by *Burkholderia* T6SS-5, where a dynamic T6SS-5 is present at the poles of intracellular bacteria. We next identify and characterize the PAAR5 component of the T6SS-5 apparatus, which together with trimeric VgrG5 is predicted to form the tip of the apparatus. Through targeted mutagenesis and cellular infection experiments, we demonstrate that PAAR5 is required for fusogenic activity but dispensable for the operation of T6SS-5. A screen of *Bt* transposon mutants failed to identify additional *paar* and *vgrG* loci that were critical for cell fusion. *Bp* mutants that lacked fusogenic activity but retained T6SS-5 functionality were avirulent in the hamster model of acute melioidosis. These experiments demonstrate a concordance between the fusogenic and virulence phenotypes *in vitro* and *in vivo*, supporting an essential role for fusogenic activity in virulence. This study provides a toolkit for further understanding the role of T6SS-5 and *Burkholderia-mediated* cell fusion in pathogenesis.

## RESULTS

### Dynamic T6SS-5 activity is present at the leading and lagging poles of intracellular *Burkholderia*

Our proposed model of the *Burkholderia* intracellular lifecycle and T6SS-5-mediated cell fusion is based on work from our group (12, 51) and others (26, 31, 42, 47). As shown in Fig. 1, cytoplasmic bacteria are propelled by BimA actin-polymerization or Fla2-flagellar motility against the inner face of the plasma membrane with sufficient force to promote contact with the membrane of an adjacent cell (12, 22). We hypothesize that membrane fusion is triggered by the T6SS-5 apparatus at the forward pole of the bacterium by sequentially joining the outer and inner leaflets of plasma membranes (52). To examine the validity of this model, we constructed a *B. thailandensis* E264 (*Bt*E264) reporter strain expressing fluorescently-labeled components of T6SS-5 to examine its subcellular localization. The TssB5 apparatus sheath protein was translationally fused to monomeric cherry red fluorescent protein (TssB5-mCherry2), and the ClpV5 AAA-ATPase to superfolder green fluorescent protein (ClpV5-sfGFP). Constructs were integrated into the native genomic loci by in-frame allelic exchange (53). HEK293 cells expressing transiently-transfected blue-fluorescent actin (mAzure-actin), were infected with the *Bt*E264 *tssB5::mCherry; clpV5::sfGFP* reporter strain, which was capable of escaping endosomes, multiplying, polymerizing actin and triggering cell fusion. The localization patterns of TssB-mCherry2 and ClpV5-sfGFP were analyzed for clues on the behavior of T6SS-5.

**Figure 1.**
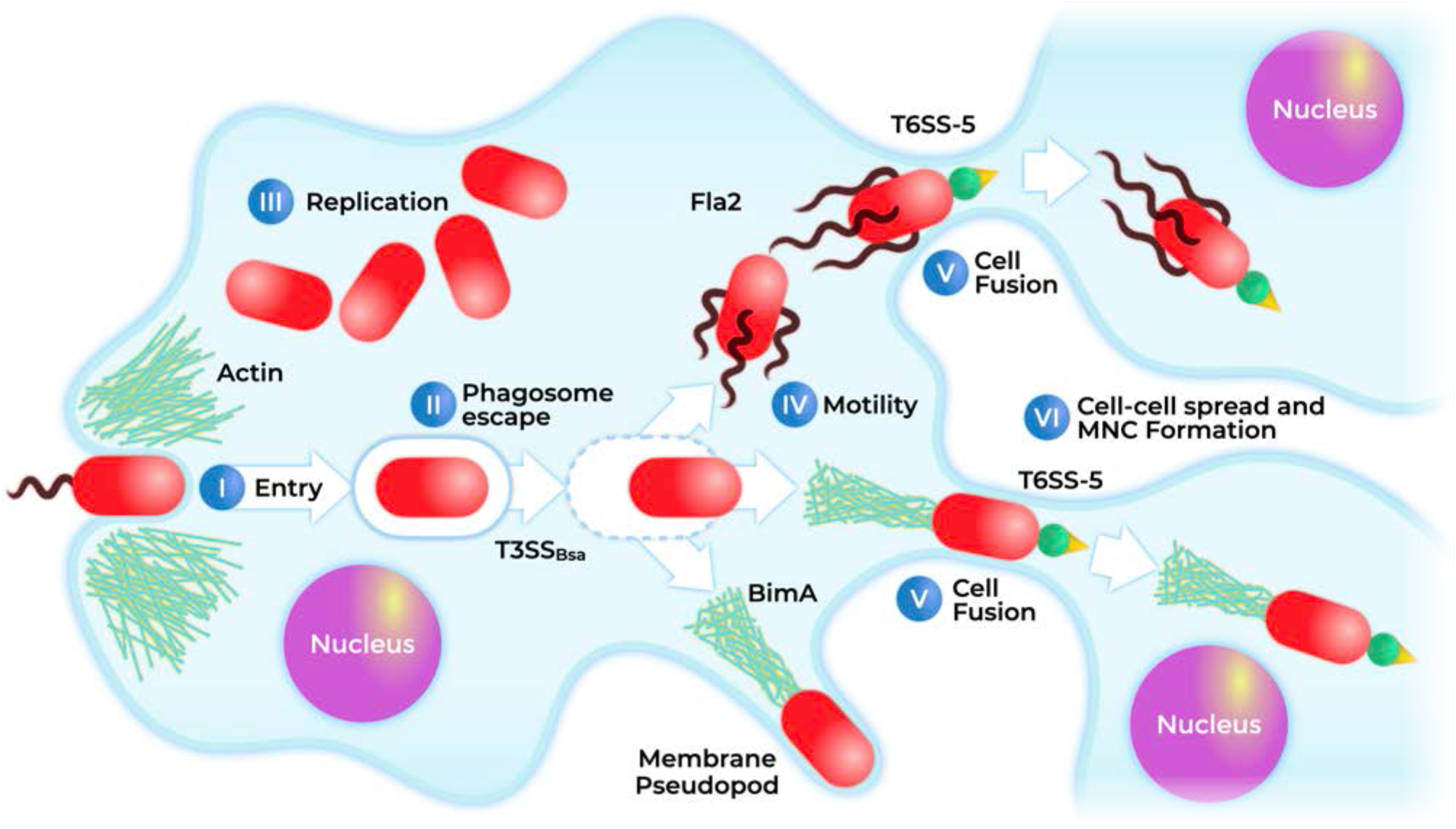
Model for intracellular lifecycle and *Burkholderia-mediated* cell fusion. *Bm, Bp* and *Bt* are facultative intracellular pathogens. Bacteria are (I) phagocytosed and the activity of T3SS_Bsa_ (II) facilitates their escape from phagosomes into the cytosol, where they (III) multiply and (IV) locomote by flagellar (Fla2) or actin-polymerization (BimA) mediated motility, either of which can propel them into the plasma membrane. Fusion with adjacent cells (V) is catalyzed by the activity of a polarly localized T6SS-5 organelle, creating a portal for direct transmission into a neighboring cell cytosol (VI), bypassing the need for additional rounds of escape from phagosomes.

A ‘loaded’ T6SS is denoted by multimeric TssB5 sheath structures. After ‘firing’, apparatus disassembly and component recycling are correlated with the presence of the ClpV5 AAA ATPase. As shown in Fig. 2, fluorescent foci of TssB-mCherry2 and ClpV5-sfGFP are prominent at bacterial poles, with smaller examples along the length of the cell. Foci containing either TssB-mCherry2 or ClpV5-sfGFP were observed, although colocalization more prevalent, indicated by yellow in the merged images (Fig. 2A-C). To quantitatively assess localization, we performed a meta-analysis on a large number (n=289) of intracellular bacteria. Single bacterial cells at the tips of actin protrusions were chosen for analysis. Dual, non-septate bacterial cells were also observed, but excluded (Fig. 2A). For single bacteria, the fluorescence signal intensities for the mAzure-actin, TssB-mCherry2 and ClpV5-sfGFP channels were recorded from the lagging to the leading pole along the center axis of the bacterium. Fluorescence line-intensity profiles for each channel were plotted relative to their location along the cell axis.

**Figure 2.**
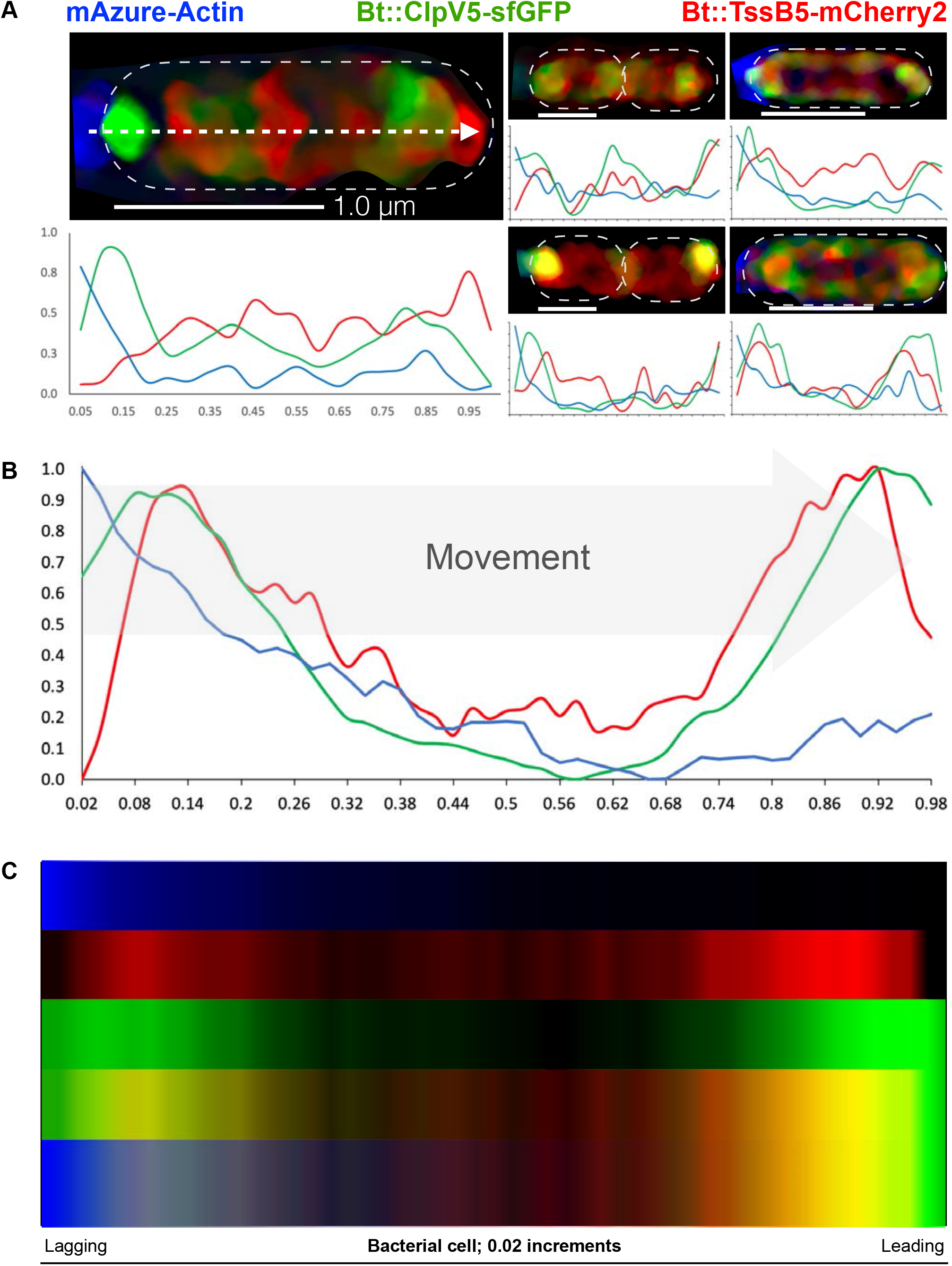
Dynamic T6SS-5 organelles are predominant at bacterial poles. (**A).** Fluorescence localization experiments. The localization and dynamics of T6SS-5 were visualized using a *Bt*E264 strain expressing fluorescent components of the T6SS-5 apparatus (29), and examples of dual non-septate and single bacteria are shown. Monomeric cherry red fluorescent protein (mCherry) was translationally fused to TssB5, a component of the apparatus sheath, to mark loaded injectisomes. Locations corresponding to sheath disassembly are denoted by the ClpV5 AAA-ATPase fused to superfolder GFP (sfGFP). These constructs were inserted into the genome of *Bt*E264, and this reporter strain was used to infect HEK293 cells expressing monomeric blue-fluorescent protein fused to cellular actin (mAzure-actin). Cells were prepared for fluorescence microscopy 18 h after infection. Micrographs and the corresponding charts of fluorescence intensity (Y axis) over cell length (X axis). mAzure-Actin (blue), TssB5-mCherry (red), and ClpV5-sfGFP (green). Fluorescence intensities of mAzure-Actin (blue), TssB5-mCherry (red), and ClpV5-sfGFP (green) were measured along the bacterial cell centerline as shown at the upper left **(B).** Fluorescence intensities for a population of intracellular *Bt*E264 reporter strain bacteria (n=289) were measured along the bacterial cell centerline and represented in chart format. BimA-dependent actin polymerization occurs at the lagging pole of the bacterium (left) as expected, as indicated by an intense blue signal. Dynamic T6SS organelles are localized at the lagging (left) and leading (right) bacterial poles denoted by discrete regions of fluorescence, where red and green correspond to loaded and fired injectisomes, respectively. Loci exhibiting yellow fluorescence, produced by spatial overlap of red and green, were common implying that T6SS-5 dynamics are high at the leading and lagging poles. The polar localization and dynamic behavior of T6SS-5 are consistent with our current model of T6SS-5 operation during cell fusion by *Burkholderia*. **(C).** Fluorescence spectrograph of fluorescent channels. Top to bottom: mAzure-Actin (blue), TssB5-mCherry (red), ClpV5-sfGFP (green), merge of green and red, merge of blue, green and red. For **B** and **C**, the two tailed t-test was used to compare the means of polar fluorescence maxima for the TssB5-mCherry and ClpV5-sfGFP channels at the leading vs the lagging cell pole, where no significant difference was found (P>0.05). The 289 bacteria chosen for this analysis encompassed 9 datasets gathered over 4 independent experiments.

The results of the meta-analysis and fluorescence spectrograph (Fig. 2B-C) provide critical insight into the operation of T6SS-5. As expected, the lagging bacterial pole is defined by a strong signal for mAzure-actin with a maximum intensity score (MIS) of 1.0, denoting the location of BimA-mediated actin polymerization. The TssB-mCherry2 sheath component exhibited an intense signal at the leading bacterial pole denoting it as a favored site for assembly and loading of T6SS-5 organelles. A prominent signal for ClpV5-sfGFP was seen to colocalize with TssB-mCherry2 at the leading pole, signifying the location of ‘fired’ T6SSs undergoing recycling. These results show that T6SS-5 behavior is highly dynamic at the leading bacterial pole, consistent with our hypothetical model and proposed role in membrane fusion (Fig. 1). Interestingly, intense signals for TssB5-mCherry and ClpV5-sfGFP were also observed at the lagging bacterial pole, indicating that it too contains dynamic T6SS-5 organelles. The difference between the red and green fluorescence maxima at the leading vs. lagging pole was not statistically significant (P>0.05 for TssB5-mCherry and ClpV5-sfGFP). In light of this observation, T6SS-5 may be performing uncharacterized functions or additional roles at the lagging pole that are distinct from its ability to trigger fusion. Understanding the implications will require further investigation.

### PAAR5 is essential for cell fusion but dispensable for the functionality of T6SS-5

In light of current T6SS structural models, we wished to further investigate the significance of polarly-localized T6SS-5 activity. We sought to identify the PAAR tip component of T6SS-5 and other PAAR homologs in *Bp, Bm* and *Bt*. The genome of *Bp* strain 1026b (*Bp*1026b; NC_017831.1, NC_017831.2) was queried using the NCBI Conserved Domain Retrieval Tool (CDART) for loci encoding the canonical DUF4150 domain of PAAR-like proteins (Pfam PF13665 and PF05488). The search returned 12 PAAR candidates for *Bp*1026b (Table 1), which were used to optimize the Delta-BLAST algorithm for PAAR proteins in other Bpc spp. We identified 9 *paar* loci in the genome of *Bp*K96243 (Table 1). One of these, *paar5*, is located within the T6SS-5 gene cluster and encodes a 130-amino acid (aa) protein with high conservation in *Bm, Bp* and *Bt* (Fig. 3A). PAAR5 is unique to membrane fusogenic Bpc spp. and diverges significantly from other *Burkholderia* PAAR analogs (Fig. 3A and Fig. S1).

**Table 1.**
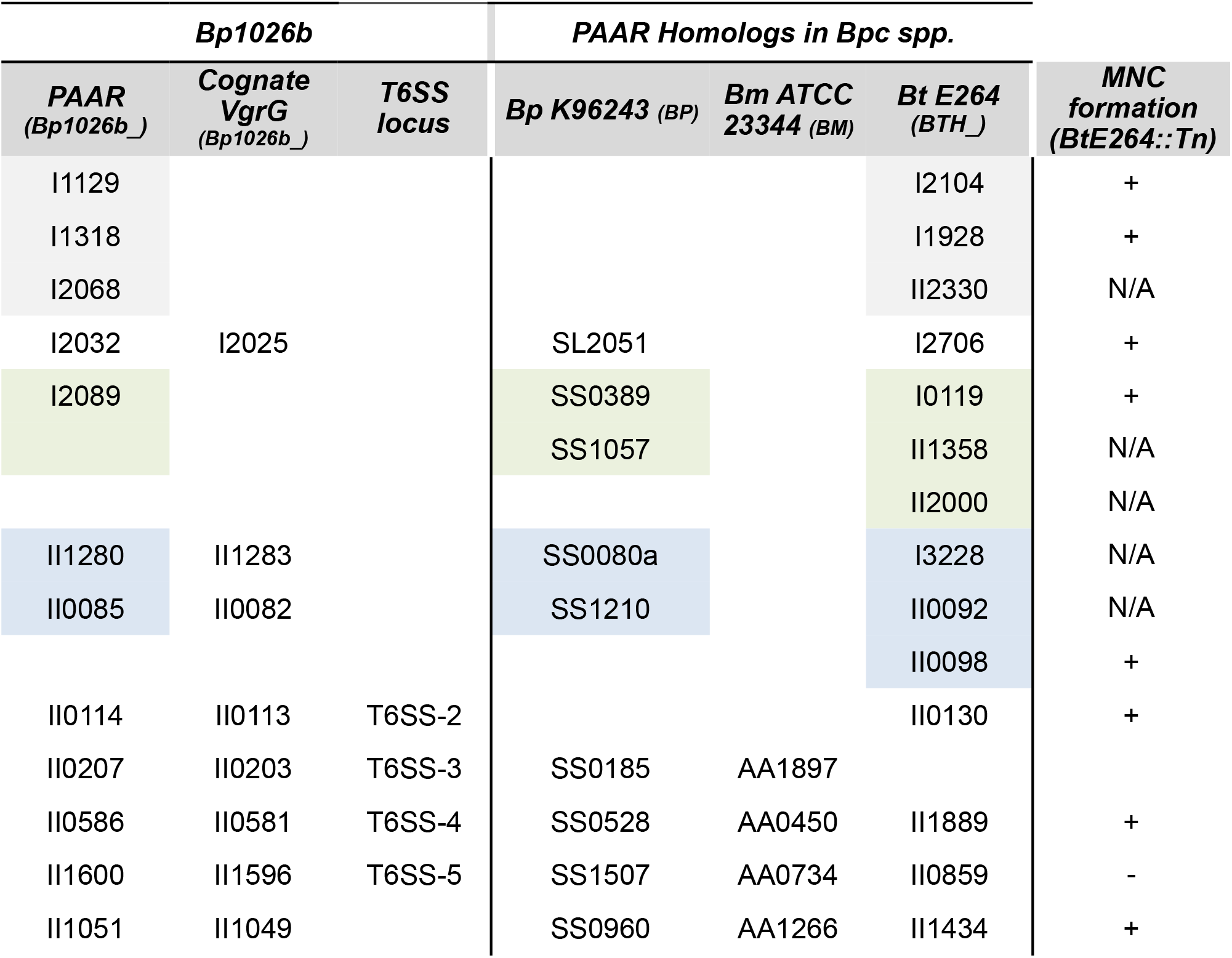
MNC formation and *paar* homologs in Bpc spp. 12 *paar* loci in *Bp*1026b were revealed by a bioinformatic search. These were used to identify homologs in *Bp*K96243, *Bm*ATCC23344 *and Bt*E264. These are grouped by probable associations with T6SS gene clusters and cognate *vgrG* loci according to chromosomal location and genomic context. LOCUS_TAG nomenclature is used (i.e. PAAR5 of *Bp*1026b is represented as Bp1026b_II1600). Color shaded boxes indicate paralogous genes possibly arising through gene duplication. Right sidebar; MNC formation phenotype observed in the transposon mutagenesis screen is indicated. +; presence of MNCs. N/A; not available in the transposon library. *paar5::Tn* was not among the *Bt*E264 library mutants but was created separately by allelic exchange mutagenesis.

**Figure 3.**
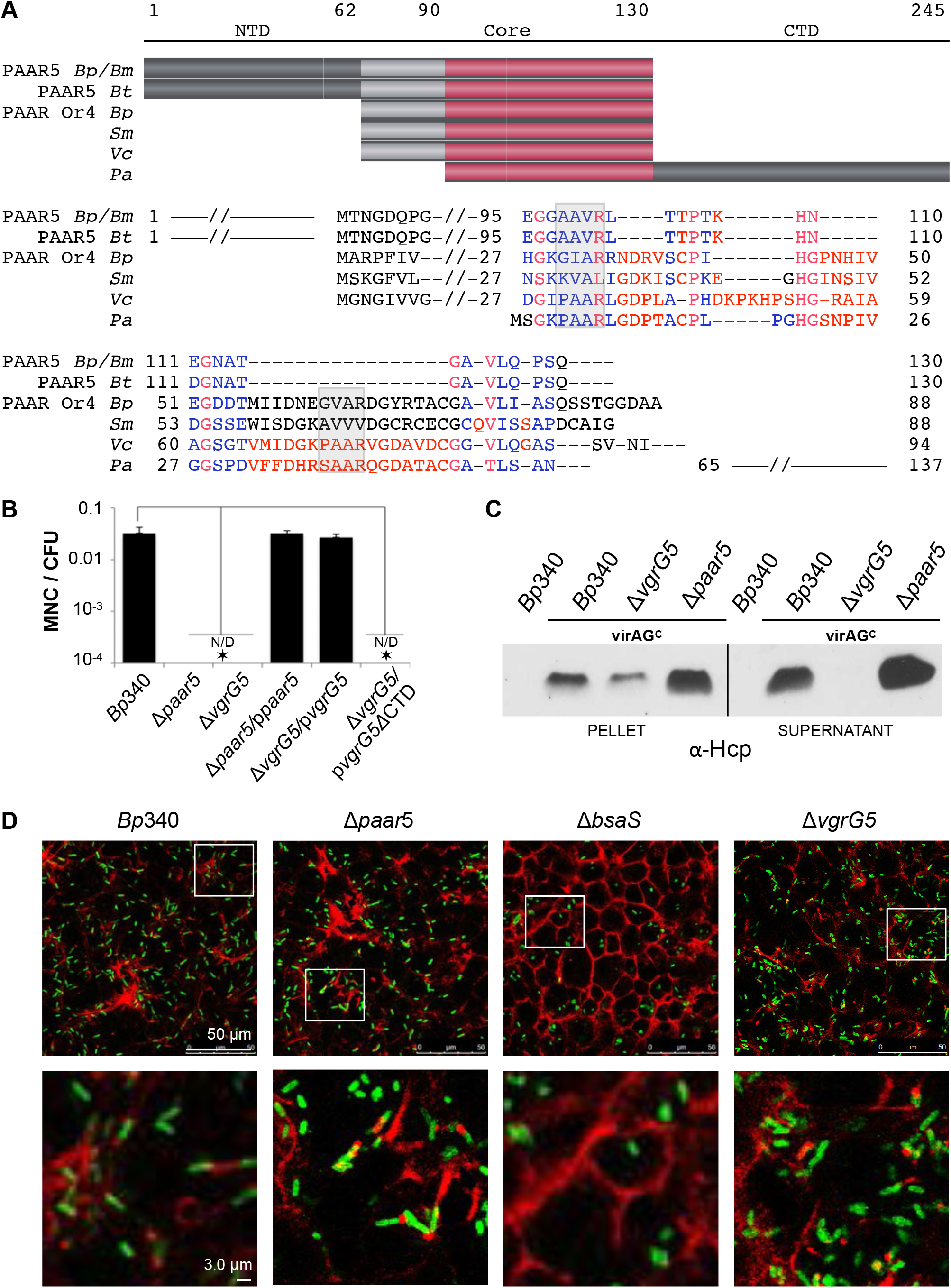
*Burkholderia* PAAR5 is required for MNC formation but dispensable for T6SS-5 operation. (**A).** PAAR5 sequence alignment from *Burkholderia pseudomallei* (*Bp*) and *Burkholderia mallei* (*Bm*), compared to orthologs in *B. thailandensis* E264 (*BtE264*), and an orphan PAAR homolog from *Bt*E264 (PAAR Or4). PAAR homologs from other species; *Vibrio cholerae* (*Vc*), Serratia marcescens (*Sc*), and Pseudomonas aeruginosa (*Pa*). Sequences (not to scale with **A.**) corresponding to highly conserved alleles in fusogenic species are highlighted in light grey. The PAAR motif comparison to *Bp*1026b is indicated in grey-shaded boxes. Functionally conserved residues; blue and red. **(B).** Assay for MNC induction efficiency per colony forming unit (MNC/CFU) with *Bp*340 and mutant derivative strains. HEK293 cells were infected with *Bp*340, Δ*vgrG5* or Δ*paar5* mutants, with and without native *paar5* alleles expressed *in trans*. MNCs were analyzed at 20 h post infection. Values represent the mean ± SD of a minimum of 3 independent experiments (* P<0.005). N/D; not detected **(C)**. Hcp secretion assay in *Bp*340 and mutant derivative strains. Western blot analysis of bacterial supernatant or pellet fractions from *Bp*340 control (WT), *vgrG5* and *paar5* mutants, and *Bp*340 *virAG* constitutive (*virAG^c^*) derivatives. Blots were probed with antibody against *B. pseudomallei* Hcp5 (α-Hcp). **(D)**. Deletion of *paar5* does not affect phagosome escape or actin-based motility. *Bp*340 and mutant derivative strains were fixed and stained 8 h after infection of HEK293 cells. Red; actin stained with Alexa Fluor 647-labeled phalloidin. Green; bacteria stained with a mouse monoclonal antibody to *Bp* LPS and goat anti-mouse Alexa Fluor 488. Bar; 50 μm. Right panel: Enlarged areas indicated by boxes in left panel. Δ*bsaS; Bp*340 T3SS_Bsa_ ATPase mutant; Δ*vgrG5: Bp*340 deleted for *vgrG5; Δpaar5: Bp*340 deleted for *paar5*. Bar; 3.0 μm. The Δ*paar5* and Δ*vgrG5* mutants escape from phagosomes and polymerize actin comparably to the *Bp*340 parental strain. The Δ*bsaS* T3SSBsa mutant remains trapped in phagosomes and cannot polymerize actin.

To study the role of PAAR5 in the *Burkholderia* lifecycle inside cells, we constructed an in-frame deletion mutant in *B. pseudomallei* strain *Bp*340 (*Bp*340), an antibiotic-sensitive efflux mutant derivative (Δ*amrAB* Δ*oprA*) of *Bp*1026b (54), and examined MNC formation following infection of HEK293 cells (Fig. 3B). In contrast to the abundant MNCs induced by the *Bp*340 parental strain, we observed no evidence of MNC formation for the Δ*paar5* mutant at 20 h post infection, comparable to the negative control Δ*vgrG5* strain that is incapable of cell fusion (Fig. 3B) (22). Complementation with native *paar5* or *vgrG5* alleles *in trans* fully restored fusogenic activity in the corresponding mutants. The VgrG5 C-terminal domain (CTD) deletion strain (Δ*vgrG5* / *pvgrG5*Δ*CTD*), which lacks sequences required for MNC formation, was unable to restore cell fusion as expected (22). The ability of Δ*paar5* mutants to invade cells, escape from phagocytic vesicles and polymerize actin was comparable to the parental strain (Fig. 3C), unlike the negative control Δ*bsaS* T3SS_Bsa_ mutant, which cannot escape phagosomes (12, 18). Since the Δ*paar5* mutant successfully entered and multiplied within cells, we conclude that PAAR5 is essential for T6SS-mediated cell fusion, and dispensable for prior steps of the intracellular lifecycle.

During T6SS deployment, contraction of the T6SS outer sheath and ejection of the inner tube structure releases Hcp subunits into the extracellular milieu. Detection of secreted Hcp5 in culture supernatants by Western Blot provides an *in vitro* assay for functionality of the T6SS-5 system (22, 55). Upon escape of the bacterium into the cytosol, normal activation of T6SS-5 is controlled by the VirAG two-component regulatory system (56). Since activity of T6SS-5 is minimal in laboratory medium, analysis *in vitro* was facilitated by constitutive expression of VirAG (*virAG*^c^) (22). Shown in Fig. 3D, the *Bp*340 parental strain lacked the ability to secrete Hcp5, while secretion was observed in *Bp*340 *virAG^c^* as expected. This contrasts with the *virAG*^c^ Δ*vgrG5* mutant, which produces but cannot secrete Hcp5 due to an essential role in T6SS-5 functionality (34). Conversely, production and secretion of Hcp5 by the *virAG*^c^-*paar5* mutant was comparable to the *virAG*^c^ parental strain. In summary, elimination of *paar5* abrogates MNC formation without disrupting the physical operation of the T6SS-5 apparatus. This phenotype is identical to that previously reported for strains carrying mutations in the VgrG5 C-terminal domain (*vgrG5-DCTD*) which, like PAAR5, is uniquely shared among fusogenic Bpc spp. (22).

### The PAAR5 N-terminus is a specialized domain that is critical for MNC formation

Although the sequence of PAAR5 is divergent from the structurally-characterized VCA0105 PAAR protein of *V. cholerae* (33) (Fig. 3A), analysis using the Phyre2 prediction algorithm (57) suggests that their overall structures are similar, with the characteristic pyramidal fold of PAAR-family members (Fig. 4A) (33, 40, 58). Compared to PAAR proteins of other species, PAAR5 carries a unique N-terminal extension (NTE) (aa 2-19) that is predicted to be structurally disordered and hydrophobic (hydrophilicity index < -1.0), implying that it is internally buried and minimally exposed to solvent. The PAAR5 NTE is followed by a proline-rich motif (PRM; aa 20-37) that may serve as a semi-rigid linker to the rest of the protein (Fig. 4B). The C-terminal region (aa 64-130) resembles the DUF4150 domain of PAAR proteins in other species.

**Figure 4.**
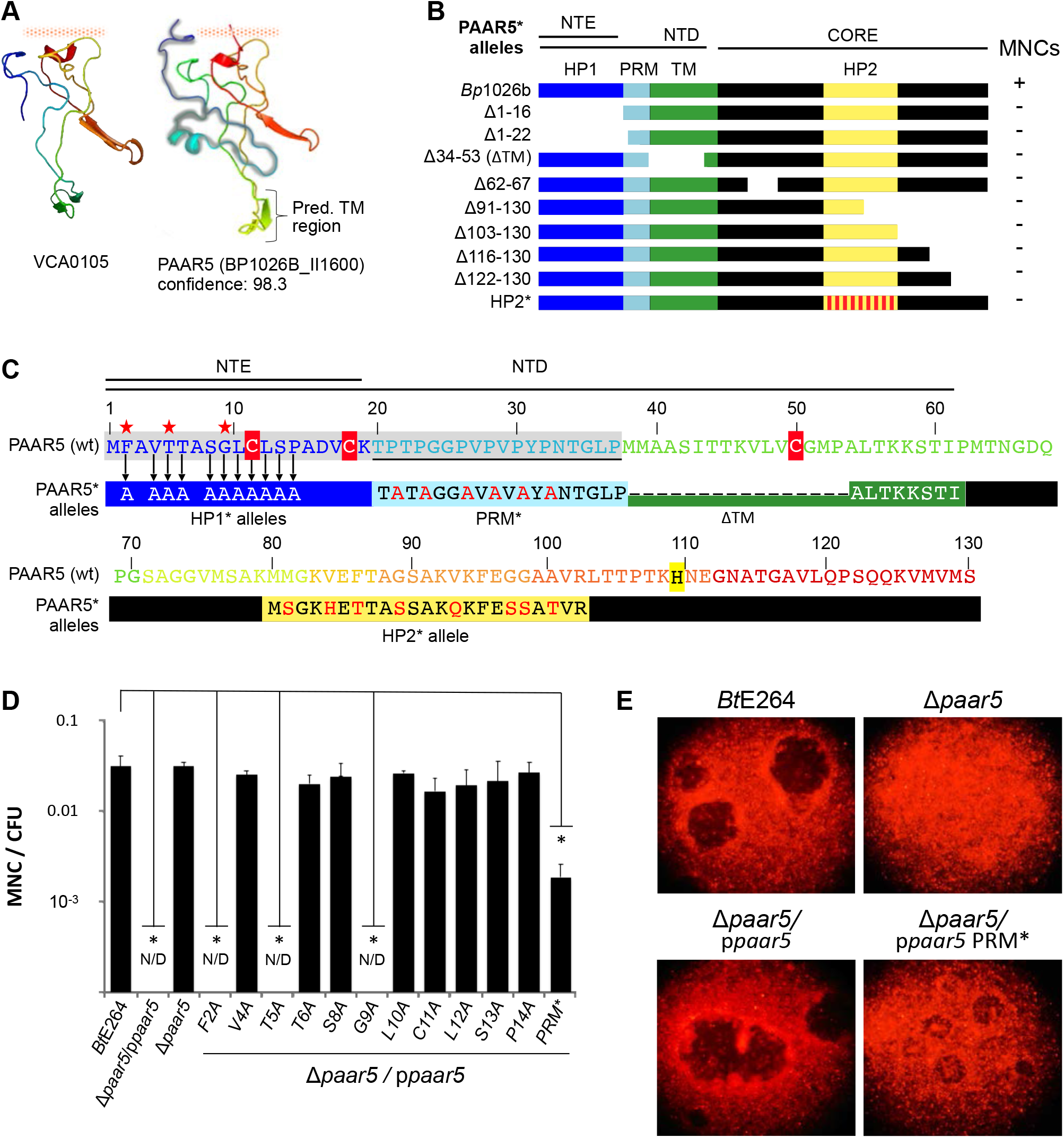
PAAR5 mutational analysis. **(A).** Phyre2 3D predictions of the PAAR protein VCA0105 from *V. cholerae* and PAAR5 of *Bp*1026b. The N-terminal extension (NTE) of PAAR5 is highlighted in grey. Confidence; Phyre2 prediction probability that PAAR5 is similar in structure to VCA0105. Red shaded bar indicates potential site of association between PAAR and VgrG. **(B).** Diagram of PAAR5 features and mutant derivatives. Sidebar; MNC formation compared to non-mutant PAAR5 of *Bt*E264. Abbreviations: N-terminal extension (NTE); N-terminal Domain (NTD); Proline Repeat Motif (PRM); Hydrophobic Patches (HP1, HP2); Putative transmembrane domain (TM). **(C).** Colored protein sequence of the PAAR5 allele in *Bt*E264. The NTE and proline rich motif (PRM) of PAAR5 are shaded in grey, and the PRM is underlined. Cysteine and histidine residues are highlighted in red and yellow respectively. Sequence alterations of mutant alleles (PAAR5*) are indicated. Arrows indicate single amino acid substitutions in mutant PAAR5 NTE alleles. Red stars indicate substitutions in the NTE that impeded MNC formation. **(D)**. MNC formation efficiency of mutant *paar*5 alleles expressed in a Δ*paar5* mutant compared to wild-type *paar5*. MNCs per colony forming unit (MNC/CFU) were determined at 20 h post infection. N/D; not detected. Values represent the mean ± SD of a minimum of 3 independent experiments (* P<0.005). **(E)**. HEK293 cells. Representative MNCs appear as dark areas in a cell monolayer 20 h post-infection with *Bp*340, Δ*paar5*, or Δ*paar5* mutant complemented with the wild-type *Bp*340 *paar5* allele (Δ*paar5* / *ppaar5*), or a proline-rich motif deletion mutant allele (Δ*paar5* / *ppaar5 PRM*\*), which reduced but did not eliminate MNC formation.

PAAR5 does not possess signal sequences that suggest periplasmic localization, although it does contain a possible transmembrane (TM) region (aa 38-63; TMpred score < -500) (59), and an additional predicted hydrophobic patch in the core of the protein (HP; aa 80-100) (Fig. 4B, C). This contrasts with the canonical *Vibrio* VCA0105 PAAR homolog which lacks hydrophobic regions. To identify sequences important for cell fusion, we constructed mutant PAAR5 alleles (PAAR5*) with terminal truncations, internal deletions, or sequence substitutions (Figs. 4B-C). These constructs were expressed in a fusion-defective Δ*paar5* deletion strain *in trans* and assessed for their ability to complement MNC formation. The Δ1-16 and Δ1-22 alleles were defective for MNC formation, lacking parts (Δ1-16) or all (Δ1-22) of the NTE, indicating a functionally critical region of PAAR5. Additional fusion-defective alleles were the ΔTM* mutation, and the allele containing substitutions in the HP2 region within the PAAR5 core motif (HP2*). Complementation with a PAAR5 allele containing proline-alanine substitutions in the proline rich motif (PRM*) impaired, but did not eliminate MNC formation, suggesting that the prolines in this region are important but not required for the function of the PAAR5 (Fig. 4C-E).

The PAAR5 NTE is predicted to extend as a disordered coil from the T6SS-capping core motif of PAAR5. To more closely examine the role of this region, we engineered eleven PAAR5 derivatives by performing alanine scanning mutagenesis within the NTE (Fig. 4C). Three individual substitutions within the first 9 aa were capable of abolishing MNC formation, while single substitutions from L10 to P14 had no measurable effect (Fig. 4D), suggesting that the first 9 aa of the NTE constitutes a specialized domain of PAAR5 with a critical role.

### Roles of additional *B. thailandensis* PAAR and VgrG proteins in cell fusion

The essential roles of PAAR5 and the VgrG5-CTD in cell fusion, along with their predicted colocalization at the tip of the T6SS apparatus (33, 36) and their dispensability for T6SS secretion activity, support a scenario where PAAR5 functions together with the VgrG5-CTD as a unique puncturing device that triggers fusion between mammalian cell membranes. In light of the numerous T6SSs encoded by fusogenic Bpc spp., we wondered if the additional T6SSs, *paar* or *vgrG* loci were contributing to cell fusion. To test this, we selected all the available *paar* and *vgrG* mutants from a curated two-allele transposon mutagenesis library in *Bt*E264 (60), and assessed their roles in MNC formation in mammalian cells (Table 1, Table, Table S2).

When compared with *Bp*1026b, the genome of *Bt*E264 lacks T6SS-3 (Table S1). Moreover, no *paar* or *vgrG* loci are linked to T6SS-1 in *Bm, Bp* or *Bt* (Table 1). T6SS-1 may share a VgrG and PAAR protein with another system or rely on unlinked ‘orphan’ VgrG or PAAR proteins. In *Bt*E264, transposon insertions in loci belonging to T6SS-1, -2, -4 and -6 did not impede MNC formation relative to the parental strain (Table S1). This includes insertions in the T6SS-linked *paar* genes (Table 1) and the five *vgrG* loci linked to these systems (Table S2). Likewise, no fusion defect was recorded for insertions in 7 of 8 orphan *vgrG* loci that are unlinked to the T6SS gene clusters (Table S2), nor for 5 of 10 orphan *paar* genes (Table 1). Mutants for the remaining 5 orphan *paar* genes were not present in the transposon library, likely not being disrupted due to their small size. While a possible role for other PAAR and VgrG proteins in cell fusion cannot be completely excluded based on these results, it is noteworthy that orthologs of these loci are lacking in *Bm*, which can efficiently trigger cell fusion with a smaller genome (5.8 Mb for *Bm vs*. 6.2 and 6.8 Mb for *Bt* and *Bp* respectively) and a markedly reduced set of *vgrG* and *paar* genes (Table 1).

Since VgrG and PAAR proteins may bind and deliver cargo effectors (33, 36), we considered the possibility that membrane fusion involves one or more cryptic effector proteins that depend on T6SS-5. Effector genes of other organisms’ T6SSs are normally located near their cognate *vgrG* and *paar* loci (61–66). In *Bm, Bp* and *Bt* loci flanking the *vgrG5* and *paar5* genes encode conserved components of the T6SS-5 apparatus; their disruption was detrimental to cell fusion, as was disruption of other structural genes within the T6SS-5 cluster (Table S2). Bioinformatic inspection of the *bim* motility and T3SS_Bsa_ gene clusters, which flank T6SS-5 but do not participate in membrane fusion (12), likewise did not reveal evidence of fusogenic T6SS-5 effector candidates. The single effector locus within the T6SS-5 gene cluster, denoted as *tssM*, encodes a ubiquitin hydrolase that is coregulated by the VirAG two component system and exported by the type II secretion pathway (26, 67). Transposon insertions in *tssM* did not eliminate cell fusion. Insertions in T3SS_Bsa_ were predictably impaired cell fusion due to their inability to escape from phagosomes (22). This contrasts with transposon insertion mutants for a phytopathogen-like T3SS cluster of *Bt*E264, partially analogous to the conserved T3SS-2 locus of *Bp* (68), which formed MNCs as efficiently as parental *Bt*E264 (Table S1). Overall, these results support the hypothesis that the additional T3SSs, T6SSs, *vgrG* and *paar* loci of *Bp* are not needed for cell fusion, and operation of the T6SS-5 injectisome itself may be sufficient. Critical effector proteins, if present, await discovery and characterization.

### Cell fusion and virulence

The essential roles of T6SS-5 in cell fusion *in vitro* and virulence *in vivo* are well established (26, 69), although their relationship remains undefined. Since most T6SS-5 mutants characterized thus far are defective in T6SS assembly and activity (22, 26), their avirulent phenotypes could reflect T6SS-dependent activities that are unrelated to the capacity to fuse infected cells. We took advantage of the ability to separate membrane fusion from the functionality of the T6SS-5 apparatus to more deeply explore the relationship between fusogenic activity and pathogenesis.

Syrian hamsters are highly sensitive to *B. pseudomallei* infection, with an LD_50_ of as few as 10 CFU (70). Groups of 5 juvenile female hamsters were inoculated via the intraperitoneal route (i.p.) with 10, 100, or 1000 CFU of parental *Bp*340 or mutant derivatives (Fig 5A). Mutants included a Δ*vgrG5* strain deficient for both T6SS-5 operation and cell fusion, along with the *vgrG5*Δ651-697 (22) and Δ*paar5* mutants, which retain apparatus functionality, as shown by their ability to export Hcp5, but are defective in cell fusion, phenocopying a Δ*vgrG5* negative control strain in cell infection assays (Fig. 3B, Fig. 5B). All mutations were created via markerless exchange with chromosome alleles.

**Figure 5.**
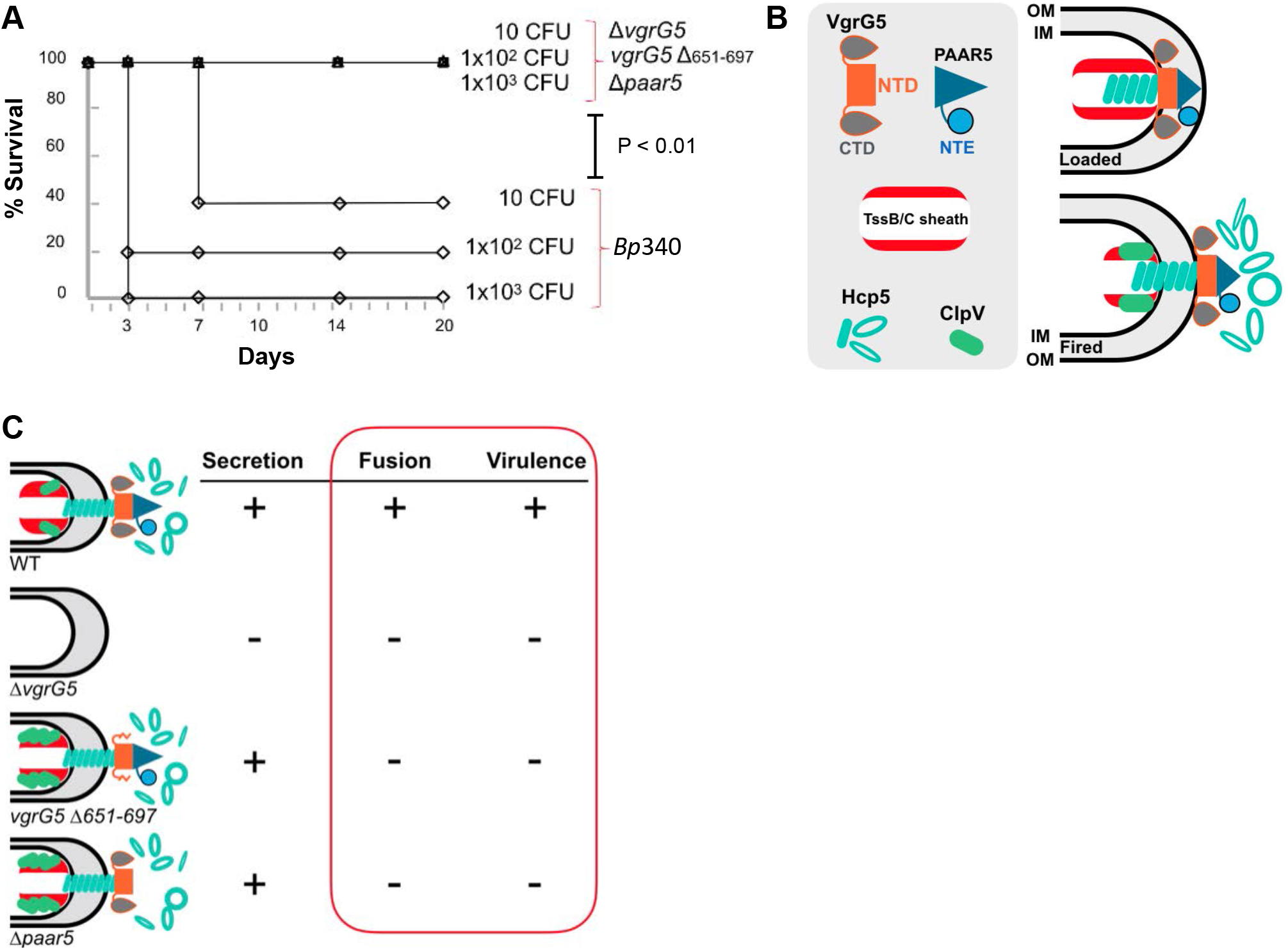
The fusogenic phenotype is correlated with virulence in animals. **(A)**. Groups of 5 female syrian hamsters were infected by the intraperitoneal route with 10, 1×10^2^ or 1×10^3^ colonies forming units (CFU) of *Bp*340 or the indicated mutant derivatives; Δ*vgrG5; Bp*340 *vgrG5* deletion strain. *vgrG5* Δ651-697; *Bp*340 mutant carrying a VgrG5 allele lacking an internal predicted 46 aa transmembrane segment in the carboxy-terminal domain (CTD). Δ*paar5; Bp*340 *paar5* deletion mutant. Animals were monitored for signs of illness for 21 days, and moribund animals were euthanized. Animals receiving mutant strains all survived through the experimental endpoint. **(B)**. T6SS-5 apparatus activity and the proposed localization of components in the loaded and fired states. Components are not to scale. The tube apparatus composed of Hcp5 subunits (light blue) and the association with the trimeric VgrG5 core structure (orange) within the extended TssB/C sheath structure (red). The VgrG5 CTD is represented by gray teardrop-shaped extensions, only two of which are shown in the diagram. The VgrG5 trimer is capped by the PAAR5 spike protein (Dark Blue) with the N-terminal extension (NTE). Apparatus firing is accompanied by contraction of the apparatus sheath and ejection of the Hcp tube and VgrG/PAAR complexes. The ClpV5 ATPase (green) localizes for disassembly of the contracted sheath and recycling of the sheath subunits. **(C).** Summary of T6SS-5 secretion, cell fusion, and virulence phenotypes. The ability to trigger cell fusion is correlated with virulence. The Δ*vgrG5* mutant (22) is defective for Hcp secretion, fusion, and is avirulent in animal infections, in contrast to wild-type (WT). The *vgrG5* Δ651-697 CTD mutation retains secretion (22) but is unable to fuse cells and is avirulent, which is phenotypically identical to the Δ*paar5* mutant.

Following infection, animals were monitored for signs of illness and euthanized if moribund. As shown in Fig. 5A, delivery of 10 colony forming units (cfu) of *Bp*340 resulted in 60% morbidity, and all animals that received 1000 cfu were moribund by day 3. The T6SS-defective Δ*vgrG5* strain was fully attenuated, with all animals surviving through the experimental endpoint. Likewise, all animals receiving the fusion-defective *vgrG5*Δ651-697 or Δ*paar5* mutant were symptom-free for the 21-day course of the experiment, comparable to those inoculated with the avirulent Δ*vgrG5* control. These results reveal a full correlation between cell fusion phenotypes *in vitro* and virulence *in vivo*, supporting fusogenic activity as a central feature of virulence.

## DISCUSSION

The ability of *Bp* to fuse mammalian cells was recognized more than 20 years ago as a unique means of cell-cell spread for a bacterial pathogen (19, 71). Recent observations demonstrate that the virulence-associated T6SS-5 performs a direct and essential role in this process (22, 26, 45, 47). While advances have been made in understanding T6SS function in other species (42), this has been challenging for fusogenic *Burkholderia*. This is due in part to the presence of multiple T6SS systems and numerous accessory VgrG and PAAR proteins with uncharacterized functions. Using protein sequence analysis, structural predictions, targeted mutagenesis, genetic screening and animal virulence studies, we conclude that cell fusion is centrally involved in *Burkholderia* virulence and is likely triggered exclusively by a component of the T6SS-5 tip complex. Mutations in PAAR5 and the VgrG5 CTD are sufficient to block fusogenic activity without affecting T6SS-5 secretion functions, and the corresponding mutants were avirulent in the highly sensitive Syrian hamster model.

A plausible mechanism for cell fusion by *Burkholderia* involves the propulsion of motile bacteria against the cell membrane, and localization of active T6SSs at the front pole of the bacterium. Consistent with this, we observed that a majority of fluorescently-labeled TssB5 sheath and ClpV5 ATPase components colocalized at the polar regions of intracellular bacteria. This denotes that the bacterial poles are sites of dynamic T6SS activity. This is consistent with the load and fire paradigm for T6SSs and with our model of *Burkholderia* cell fusion, indicating successive rounds of assembly, apparatus firing and disassembly, which to our knowledge has not been reported previously. Recently, polar localization of ClpV5 in *Bt*E264 has been shown to be independent of other components of T6SS-5, as a ClpV5-sfGFP fusion protein was observed at cell poles in a strain lacking the entire T6SS-5 gene cluster (72). Whether events occurring at the lagging bacterial pole are related to fusion of mammalian cell membranes remains to be elucidated.

By mutational decoupling of fusogenic activity from T6SS-5 secretion, we found that PAAR5 was dispensable for the system’s functionality. Studies in other species suggest that at least one PAAR protein is required for T6SS assembly and function, and several PAARs may be able to individually support the activity of a single system (33, 41). This could be the case for T6SS-5, as *Burkholderia* contains numerous unlinked, orphan *paar* loci whose expression and activities are uncharacterized. Since 5 of 12 *paar* loci were not represented in our *Bt*E264 transposon library, we were not able to directly examine their relationship to MNC formation. These loci are unlinked from *Burkholderia* T6SS-5 and have not been described in previous studies of the bacterial secretome, or as under the control of the VirAG regulon, and thus we speculate that only *paar5* may be needed for cell fusion.

We performed a sequence dissection of PAAR5 to identify features important for its unique requirement in cell fusion. A short N-terminal extension (NTE) most likely protrudes from the conserved core structure of the protein, placing PAAR5 in the group of “evolved” PAAR family members. The core structures of most PAAR proteins, including PAAR5, typically contain 2 to 4 cysteine and/or histidine residues that spatially position a divalent metal ion (Zn^2+^, Mg^2+^, Fe^2+^) near the PAAR protein pyramidal vertex, which is thought to enhance structural stability and target cell puncturing (33, 41). PAAR5 contains three cysteine residues; C11, C18 and C50, any of which might coordinate a metal ion with H109. Often, cysteine residues are implicated in structural stability via the formation disulfide bridges, however alanine substitution of C11 (C11A), which is conserved in fusogenic Bpc spp., had no measurable effect on cell fusion implying a non-structural role, possibly in mediating protein-protein interactions. It has been shown that the N- and C-terminal regions of PAAR proteins can be specialized effector domains or adaptors for their cognate VgrGs or for cargo effectors (36, 41, 73, 74). One of the most relevant examples is *Francisella* IglG, which contains an N-terminal extension that mediates interactions with IglF, a cargo effector required for phagosome escape and intracellular survival (36). Based on this, the NTE may bestow a unique activity to PAAR5, or facilitate protein-protein interactions that are important for cell fusion.

From our analysis of transposon mutants in *Bt*E264, we did not find evidence that insertion mutants corresponding to the additional T6SSs, *paar* and *vgrG* loci contribute to cell fusion. Effector loci would be expected to be genomically-linked to T6SS-5 or coregulated under the control of VirAG (27, 48, 75, 76). Bioinformatic inspection of T6SS-5 and flanking genomic regions did not produce evidence of loci that may reasonably function as effectors. However, the screening experiments did confirm our earlier finding that either the *bim* actin motility or *fla2* flagellar gene clusters will suffice for providing motility as the driving force for cell fusion. Deeper transcriptomic analysis of the VirAG regulon should be helpful for identifying such effectors, if present. Microarray-based characterization of the VirAG regulon in *Bm* has been previously performed, and did not reveal gene loci that would be recognizable effector candidates (26).

*Bm* contains the fewest genes of all fusogenic Bpc spp., and all genes in *Bm* are represented in *Bp* (2). *Bt*E264 insertion mutants for all but one of the *Bm paar* orthologs (BMAA1897) were present in our transposon library, and these were able to trigger MNC formation in cells. The unrepresented *paar* locus is linked to T6SS-3, which is absent in *Bt*E264. Thus at least in *Bt*E264, an ortholog of BMAA1897 is not needed for cell fusion, since *Bt*E264 is competent to fuse cells without it. Since *Bm* efficiently fuses cells without novel genes, we may be able to exclude loci that are not shared among *Bm, Bp* and *Bt* as requirements for cell fusion, although this requires experimental verification. Such a finding would be consistent with previous studies that have not identified additional T6SS-5 effectors, including RNA-seq analysis of the VirAG- and BsaN regulons of *Bp* (75) (Y.H. Gann, personal communication and unpublished data), and a study where Tn-seq was utilized to identify virulence determinants for respiratory melioidosis (76). Moreover, mass spectrometry analysis of the VirAG-dependent secretome of *Bt* suggests that VgrG5 and Hcp5 are the only proteins secreted in a T6SS-5- dependent manner (47). Therefore, it is feasible that T6SS-5 can mediate membrane fusion without the aid of effectors and other gene products.

The Syrian hamster model of acute melioidosis was used to ascertain the relative virulence potentials of fusogenic and T6SS-defective mutants. Hamsters are highly sensitive to *Bp* infection, making this model suitable for identifying essential factors for pathogenesis (70, 77–79). Mutants that are defective for cell fusion only (Δ*paar5* or *vgrG5*Δ651-697 mutants) proved to be avirulent, resembling the Δ*vgrG5* control strain that is defective for fusion but also defective for T6SS-5 functionality. Our results reveal a concordance between fusogenic and virulence phenotypes, both of which rely on the activity of T6SS-5. Despite their strong association, the relationship between fusogenic activity and virulence has not been directly addressed. MNCs have been observed in animal and patient tissues (24, 80), and cell fusion is presumed to occur during infection, but to our knowledge, a cause-and-effect relationship has not been established. An attractive possibility is that fusogenic activity and subsequent cell-cell spread may protect the bacterium from attack by host immune responses. Our panel of T6SS-5 and fusion-defective mutants will be useful for investigating these aspects of the *Burkholderia* disease processes, and for investigating virulence differentials associated with organ and tissue involvement, or for various infection routes and host responses. To understand these features of infection, less fulminant animal models, such as BALB/c mice, will be more desirable (47, 81).

Although a functional T6SS-5 is required for virulence in mammals, the factors that maintain conservation of T6SS-5 across virulent and avirulent species are unclear. *Bp* is an accidental pathogen of mammals, and almost all infections are acquired from the environment (10). We hypothesize that the original targets of *Bp* virulence systems, such as T6SS-5 and T3SS_Bsa_, may be bacterivorous protozoa that cohabit in environmental soil and water. Discerning the ecological roles of these systems will be enlightening for understanding virulence evolution in *Burkholderia*, and will necessitate studies with environmental species.

## MATERIALS AND METHODS

### Bacterial strains and mutant construction

*Bp*340 (*Bp*1026b Δ*amrAB-oprA*) (82) and derivative strains were grown in L-broth without NaCl (LB-NS). Kanamycin or Zeocin was added at final concentrations of 100 or 500 μg/ml as required. Strains constitutively expressing *virA* and *virG* were constructed as described (22). The *Bp*340 *virAG* genes (*Bp1026b_1587-89*) were inserted downstream from the S12 ribosomal subunit promoter in a mini-Tn*7*-Zeo transposon (83). Following insertion into the *Bp* genome, clones were selected for resistance to 2.0 mg/mL Zeocin. In frame deletions of the *paar5* (BP1026B_II1600), and allele replacements with *tssB-mCherry2* and *clpV5-sfGFP* were constructed using the *pheS*\* negative selection marker on M9 agar containing 0.1% cholorophenylalanine (C-Phe) (84). Complementation of mutants was performed using derivatives of the pBBR1-MCS2 broad host-range plasmid (85) containing the *nptII* kanamycin resistance gene. Internal deletions and truncated versions of *paar5* were obtained after PCR amplification and verified by Sanger sequencing. PCR primers are listed in Table S3.

### Cellular staining and fluorescence microscopy

HEK293 cells expressing transiently-transfected actin fused to monomeric azure-fluorescent protein (mAzure-Actin; Addgene) were grown on coverslips in 12-well plates at 2×10^5^ cells/well and infected with *Bp*340, *Bt*E264 or mutants at a MOI of 10. After 1 h of infection, extracellular bacteria were killed by adding gentamicin (100 μg/ml; *Bp*340) or kanamycin (1.0 mg/mL, *Bp*1026b and *Bt*E264), and cells were prepared for microscopy as previously described (12). Antibodies were used at the following dilutions: mouse anti-*B. pseudomallei* antiserum at 1:1000, Alexa-Fluor 488- labeled phalloidin and rabbit Alexa-Fluor 633 labeled mouse antiserum (Molecular Probes) at 1:200. Permanent mounts were created using ProLong Gold (Invitrogen). Fluorescence imaging was performed using a Leica SP5-II-AOBS confocal microscope. Image processing was performed with Leica’s LAS-AF software or Adobe Photoshop.

### Localization analysis of fluorescent T6SS-5 components TssB5-mCherry2 and ClpV5-sfGFP

Cell culture and infection with the *Bt*E264 T6SS reporter strain was performed as described above. Fixation with paraformaldyde, mounting on slides was performed as described above. Microscopy was performed as described above using sequential confocal scanning with 405/488/594 nm wavelength excitation lasers for the mAzure, sfGFP and mCherry2 imaging channels, respectively. Pixel intensities for the separate fluorescence channels were recorded along the axial centerline of individual bacteria (n=289) and averaged over 2.0% (0.02) increments of the cell length to standardize the analysis among the population of bacterial cells. The lagging pole of the bacterium was defined as the location juxtaposed with an intense signal for mAzure-actin; the site of BimA-mediated actin polymerization. Data sets were de-noised by subtraction of signal background (F_min_) from signal intensity (F) according to F – F_min_ / F. The normalized fluorescence intensity (F_N_) was determined by F_N_ = F + (1.00 - F). . Line plots were constructed using the average intensities for each fluorescence channel per incremental distance from the lagging pole of a bacterium. Computational analysis was performed by MS Excel and Google Sheets. Graphs were generated and statistics were performed on the entire data set for each fluorescence channel using Microsoft Excel. Color spectrographs representing intensity vs. position were constructed in Adobe Illustrator. De-noised fluorescence intensities for mAzure, mCherry and sfGFP were pseudocolored blue, red or green respectively, and their values assigned along the length of the bacterial cell divided into 50 even segments (2% of total length).

### Transposon insertion mutant library screening procedures

A transposon insertion mutant library in *BtE264* (60) was duplicated in 96-well plates. Individual transposon mutants and control strains were inoculated in L-broth containing 15% glycerol. For propagation of the library clones prior to screening, plates were inoculated from frozen stocks and incubated at 37°C for 48 h. 10 random transposon mutants from the duplicated library were selected and the transposon insertion verified by PCR (Fig.4).

Primer for *Tn* insertion confirmation:

Tn8: Forward: 5’-TTTATGGACAGCAAGCGAACC; Reverse: 5’-Tn23: Forward:5’-TATGTGGACGAGCTGTACAAG; Reverse: 5’-AACGACGGCCAGTGAATCCG.

High-throughput screening of the Tn mutagenesis library was performed at the UCLA Molecular Screening Shared Resource (MSSR: http://www.mssr.ucla.edu). HEK 293 cells were seeded in 384-well plates at a final concentration of 3.5×10^4^ per well using a multidrop device. Bacteria from transposon library duplicated plates were diluted at 1/300 in DMEM +10% Fetal Bovine Serum. 1 μl or 10 μL of dilution were added to the 384-well plates using automated dispenser BioMek FX (Beckman Coulter). Extracellular bacteria were then eliminated by adding antibiotic to the media 1h after infection. Laser scanning cytometry data was collected for each well using a FlexStation plate reader (Molecular Devices) and analyzed for the presence of multinucleated cells and/or plaques at 20 h post-infection (PI).

### MNC formation assays

HEK293 cell lines that stably express msRFP or eGFP were infected with *Bp*340, *Bt*E264 and derivative strains as described above. RFP/eGFP expressing cells were seeded at a final concentration of 1.8×10^6^ per well pre-coated with a 1/40 dilution of liquid Matrigel in DMEM (Becton Dickinson). Cells were infected at a MOI of 1×10^-3^ and 1 h post-infection cells were washed and remaining extracellular bacteria were killed by adding kanamycin (120 μg/ml). After 20 h, cells were fixed with PBS + 10% formalin and MNCs were observed by fluorescence microscopy. The number of MNCs formed per bacterial CFU was determined and values were reported as the mean ± SD of 3 independent experiments.

### *In vitro* type VI secretion assays

Hcp5 secretion assays were performed as previously described (22). 1 ml aliquots of bacterial culture at an OD_600_ 0.5 were centrifuged, washed, and resuspended in 400 μl of Laemmli Buffer (pellet fractions). 30ml culture supernatants were filtered through a 0.2 μm syringe-tip filter, precipitated with 10% trichloroacetic acid (TCA) and resuspended in Laemmli Buffer (supernatant fractions). Supernatant fractions were normalized according to the OD_600_ of the bacterial culture at the time of harvest and samples were loaded on a 4-15% polyacrylamide gradient gel (BioRad), subjected to SDS-PAGE, and transferred to polyvinylidene difluoride (PVDF) membranes. Detection of Hcp5 was performed as previously described (22) using anti-*Bp* Hcp (rat) and secondary antibody IgG horseradish peroxidase-labeled conjugate (Amersham).

### Phylogenetic tree and 3D structures analysis

Phylogenetic trees were generated using the program Geneious after comparison of annotated PAAR-like protein sequences of *Bp1026b* (Table 1) and their homologs in *Burkholderia* strains. Predicted 3D structures of PAAR-like proteins of *Bp*1026B and VCA0105 from *V. cholerae* were obtained after submission of protein sequences to PHYRE2 program (57). Indices of confidence for the protein to be homolog are indicated as a percentage.

### Animal study

Virulence assessment of *Bp*340 strains was performed in the Syrian hamster model. Groups of 5 female Syrian hamsters, 6-8 weeks old, were infected via the intraperitoneal route with 10, 100 or 1×10^3^ colonies forming units (CFU) of the indicated *Bp*340 derivatives and were monitored for 21 days.

### Ethics statement

Animal research at the United States Army Medical Research Institute of Infectious Diseases (USAMRIID) was conducted under an animal use protocol approved by the USAMRIID Institutional Animal Care and Use Committee (IACUC) in compliance with the Animal Welfare Act, PHS Policy, and other Federal statutes and regulations relating to animals and experiments involving animals. The facility where this research was conducted is accredited by the Association for Assessment and Accreditation of Laboratory Animal Care International (AAALACi) and adheres to principles stated in the Guide for the Care and Use of Laboratory Animals (National Research Council, 2011).

## ACKNOWLEDGMENTS

The authors declare no conflicts of interest. We thank Mary Burtnick (University of Nevada, Reno) for antibody against *B. pseudomallei* Hcp, David AuCoin (University of Nevada, Reno) for antibody against *B. pseudomallei* LPS, and Robert Damoiseaux for access to the UCLA MSSR. This work was supported by grants from the Defense Threat Reduction Agency of the Department of Defense; HDTRA1-20-1-0013 to CTF, University of Arizona Regents Technology and Research Initiative Fund (TRIF) to CTF, HDTRA1-11-1-0003, and DTRA/JSTO-CBD proposal number CBCALL12-LS1-2-0070 to DD. Opinions, interpretations, conclusions and recommendations are those of the authors and are not necessarily endorsed by the U.S. Army.

## Figure Legends

**Table S1.**
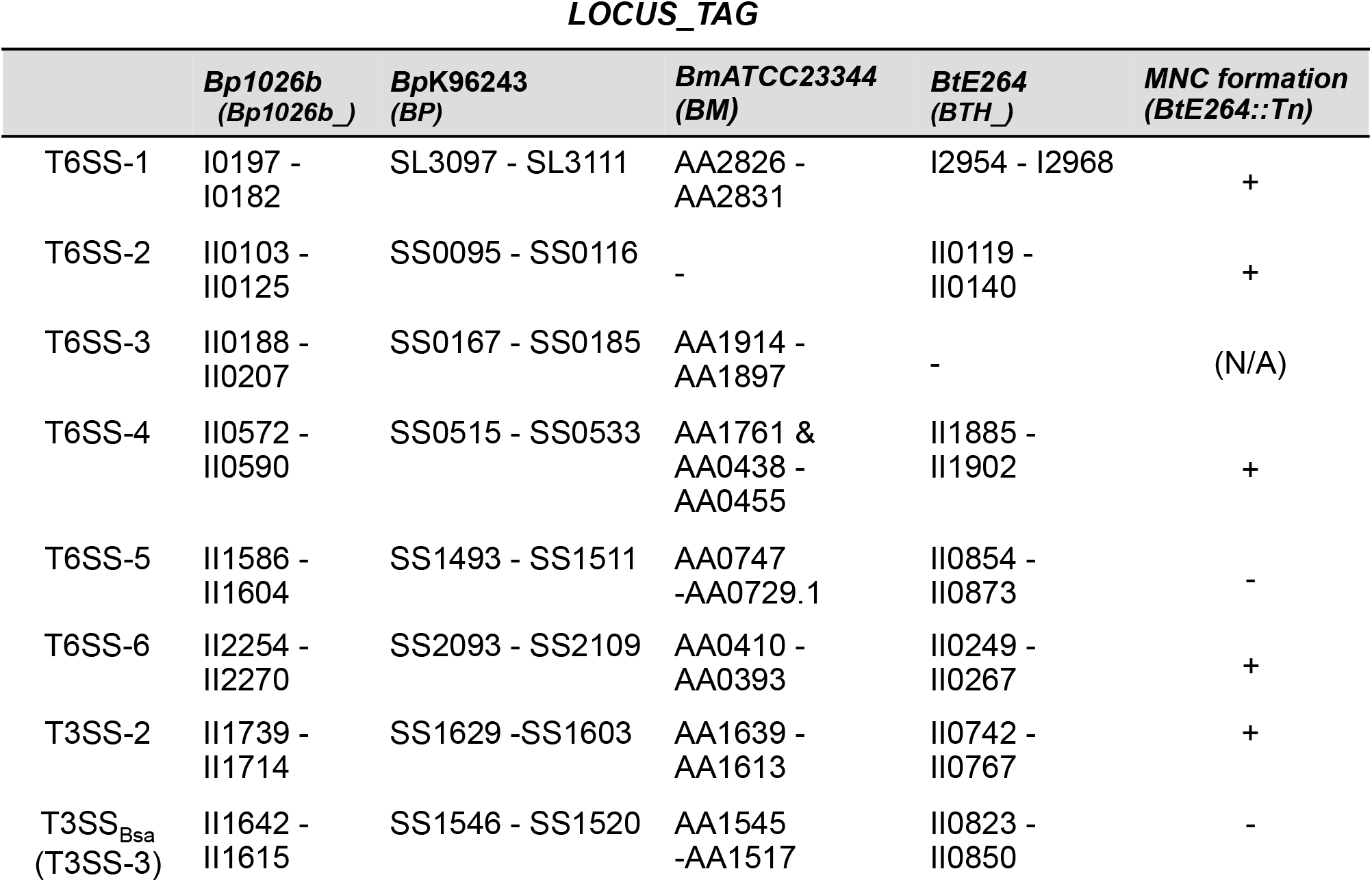
T6SS-5 is the only T6SS locus required for MNC formation. Type VI and Type III Secretion Systems (T6SS, T3SS) in fusogenic Bpc spp. The T3SS and T6SS gene clusters are denoted by LOCUS_TAG according to chromosomal location, and their distribution in other strains and species is compared with *B. pseudomallei* 1026b (*Bp*1026b). LOCUS_TAG nomenclature is used (i.e. The T6SS-5 gene cluster of *Bp*1026b is represented as Bp1026b_ II1586 - II1604). Depending on the genome annotation nomenclature, gene order in clusters may be inverted relative to others. Orthologous gene clusters in B. pseudomallei K96243 (*Bp*K96243), *B. mallei* ATCC23344 (*Bm*) and *B. thailandensis* E264 (*Bt*E264) are indicated, where present. Right sidebar; MNC formation results for *Bt*E264 transposon library mutants carrying insertions in loci encoding structural components of the T3SS and T6SS gene clusters are indicated by “+” or “–”, as positive or negative for MNC formation in the screening assay. N/A; not applicable, as the T6SS-3 ortholog is absent from *Bt*E264. Note that *Bm* lacks an ortholog of *Bp* T6SS-2.

|  | ***Bp1026b***<br>**(*Bp1026b_*)** | ***BpK96243***<br>**(*BP*)** | ***BmATCC23344***<br>**(*BM*)** | ***BtE264***<br>**(*BTH_*)** | ***MNC formation***<br>**(*BtE264::Tn*)** |
| --- | --- | --- | --- | --- | --- |
| T6SS-1 | I0197 -<br>I0182 | SL3097 - SL3111 | AA2826 -<br>AA2831 | I2954 - I2968 | + |
| T6SS-2 | II0103 -<br>II0125 | SS0095 - SS0116 | - | II0119 -<br>II0140 | + |
| T6SS-3 | II0188 -<br>II0207 | SS0167 - SS0185 | AA1914 -<br>AA1897 | - | (N/A) |
| T6SS-4 | II0572 -<br>II0590 | SS0515 - SS0533 | AA1761 &<br>AA0438 -<br>AA0455 | II1885 -<br>II1902 | + |
| T6SS-5 | II1586 -<br>II1604 | SS1493 - SS1511 | AA0747<br>-AA0729.1 | II0854 -<br>II0873 | - |
| T6SS-6 | II2254 -<br>II2270 | SS2093 - SS2109 | AA0410 -<br>AA0393 | II0249 -<br>II0267 | + |
| T3SS-2 | II1739 -<br>II1714 | SS1629 -SS1603 | AA1639 -<br>AA1613 | II0742 -<br>II0767 | + |
| T3SS<sub>Bsa</sub><br>(T3SS-3) | II1642 -<br>II1615 | SS1546 - SS1520 | AA1545<br>-AA1517 | II0823 -<br>II0850 | - |

**Table S2.**
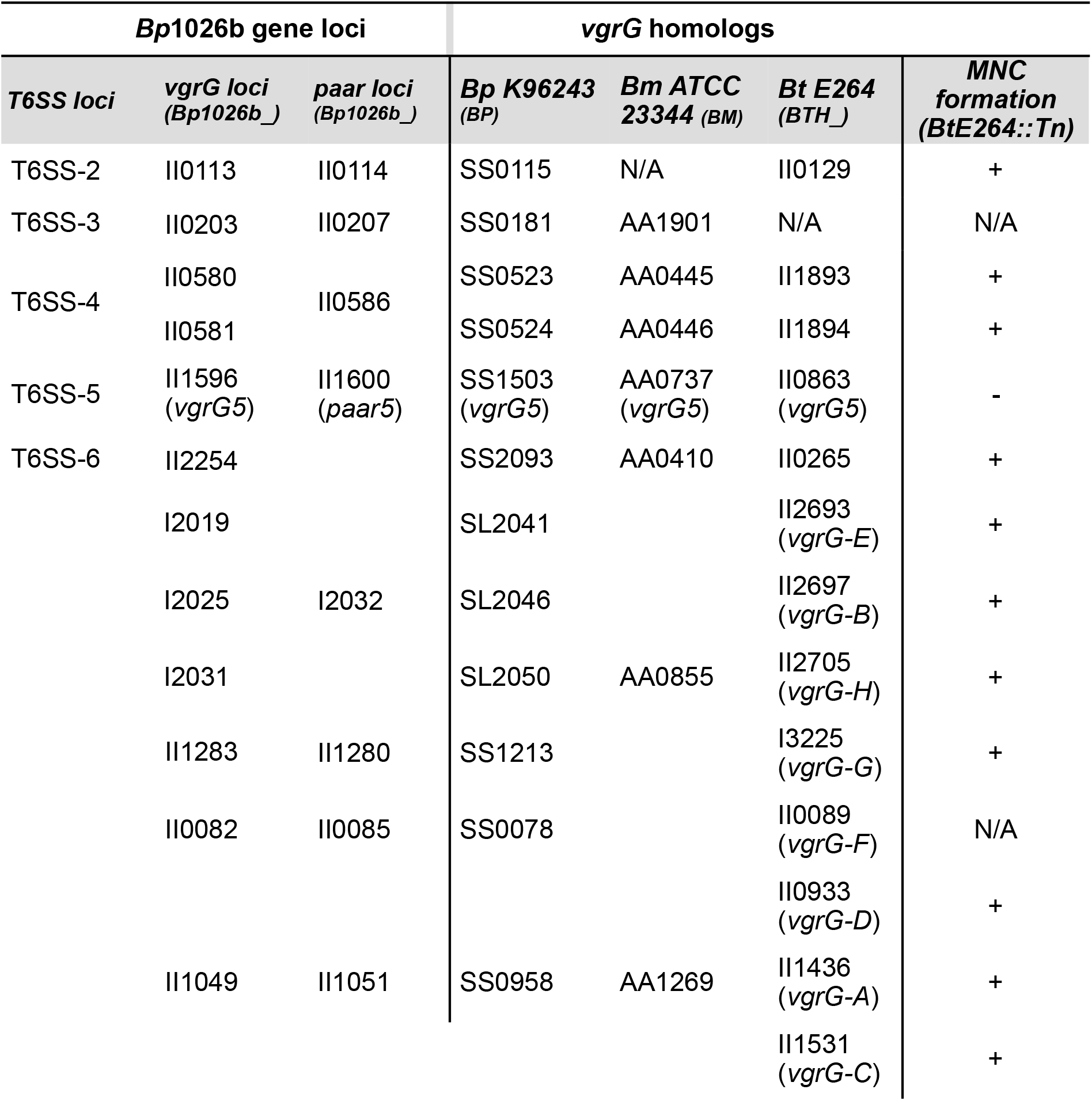
MNC formation and *vgrG* homologs in Bpc spp. 12 *vgrG* loci were revealed by a bioinformatic search using *B. pseudomallei* 1026b (*Bp*1026b). These are grouped by likely associations with their cognate T6SS and *paar* loci, where present, according to chromosomal location and genomic context. LOCUS_TAG nomenclature is used (i.e. *vgrG5* of *Bp*1026b is represented as Bp1026b_ II1596). Loci encoding VgrG homologs in *Bp*K96243, *Bm*ATCC23344 and *Bt*E264 are listed. MNC formation phenotype in *Bt*E264 mutants. +; presence of MNCs; N/A: no corresponding transposon mutant present in library. Note that *Bt*E264 lacks an ortholog of *Bp* T6SS-3 and the corresponding linked *vgrG* and *paar* loci. T6SS-1 is not shown since it lacks linked *vgrG* and *paar* genes in *Bm, Bp* and *Bt*, likely relying on orphan *paar* and *vgrG* loci for function.

**Table S3.**
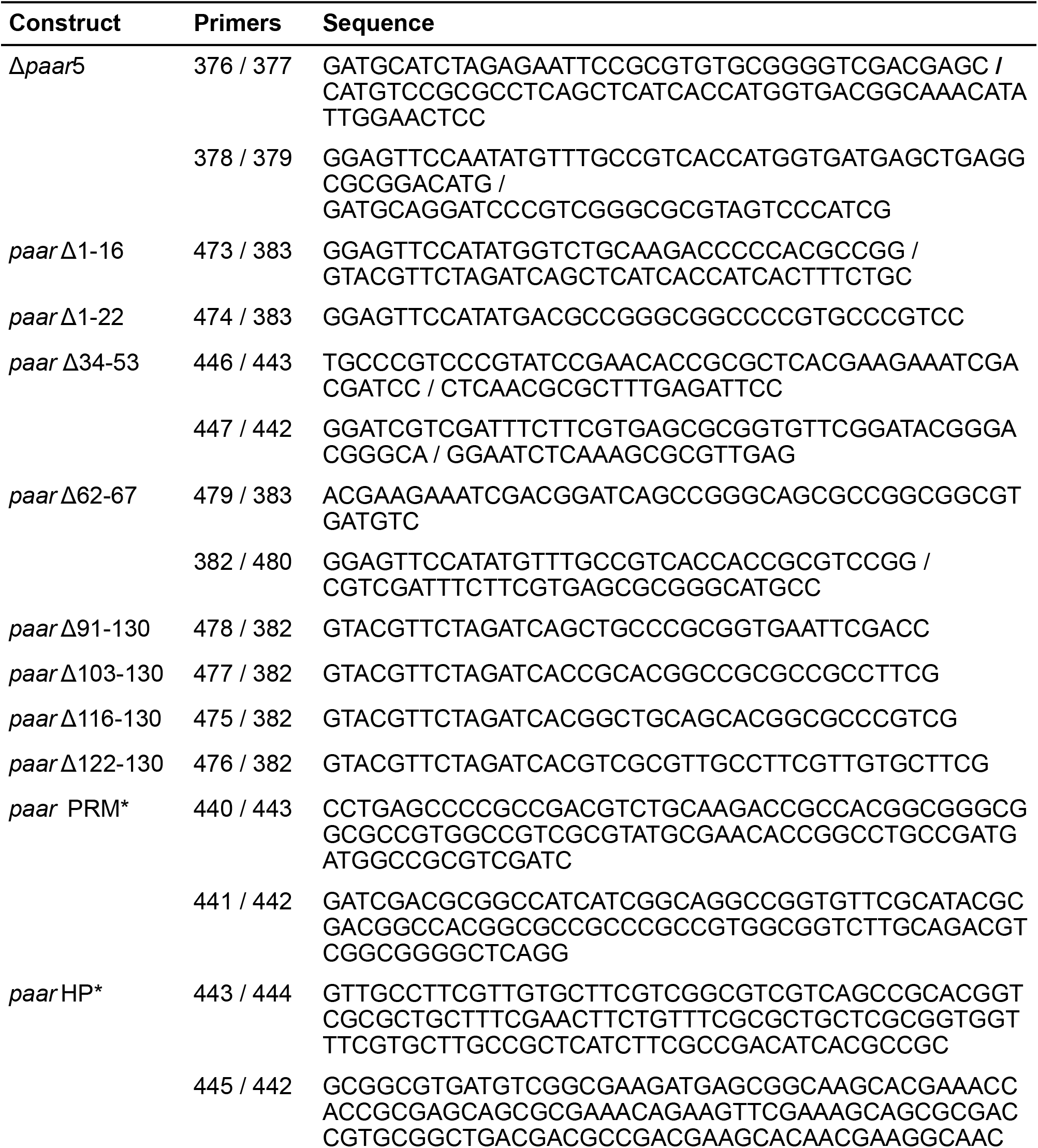
PCR primers used in this study.

**Figure S1.**
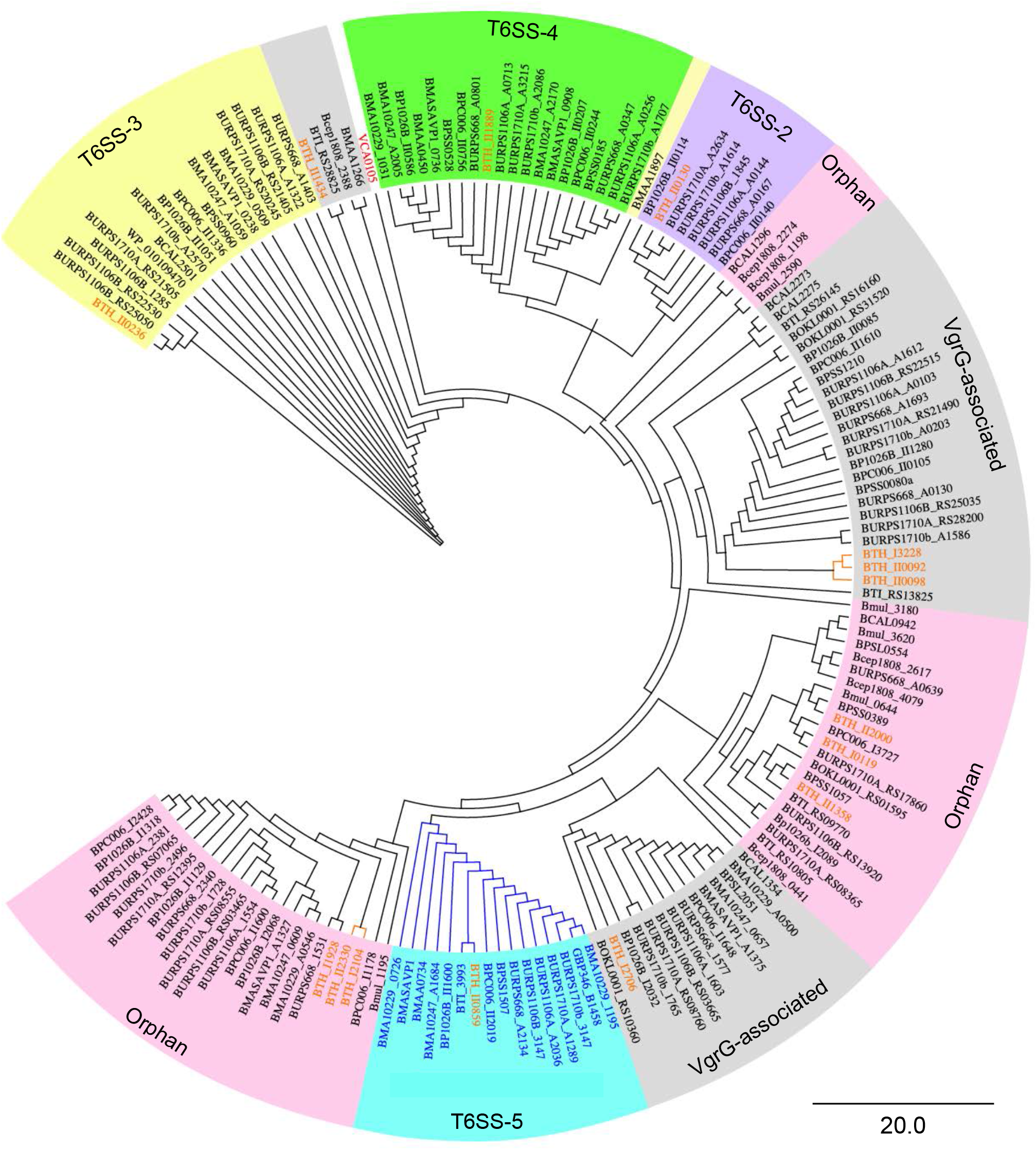
Phylogeny of PAAR proteins in Bpc spp., including *B. mallei, B. oklahomensis, B. pseudomallei*, and *B. thailandensis*. The VCA0105 PAAR protein from *V. cholerae* is shown in bold red. PAAR proteins in Bpc spp. are grouped by status; linked to a T6SS system, orphan or VgrG-associated as indicated by color shading. The diagram is rooted using PAAR proteins encoded by *Bt*E264 (red).

